# Uncertainty Shapes Neural Dynamics in Motor Cortex During Reaching

**DOI:** 10.1101/2025.10.14.682390

**Authors:** Tapas J Arakeri, Jean M Dill, Katalin M Gothard, Andrew J Fuglevand

**Author notes:** Correspondence: Andrew J. Fuglevand, Ph.D.

## Abstract

Voluntary reaching movements are often made with incomplete information about the movement goal, which may require the brain to flexibly adjust motor plans and ongoing movements. To examine how uncertainty about a reach target influences neural preparation and execution, we recorded activity from dorsal premotor (PMd) and primary motor (M1) cortices of a rhesus macaque trained to reach to one of two potential targets. In the task, a timing cue flashed three times in succession. Potential targets were displayed at the time of the first flash and the monkey needed to initiate the movement almost simultaneously with the third flash. Colorings of potential targets indicated the probability that the target would be at one location or the other, thereby inducing varying levels of uncertainty about the final target location. On half the trials, the final target was displayed so late that the monkey had to ‘guess’ the target location, based on cues provided by the potential targets. While the monkey performed this task, we recorded 165 neurons from PMd and 37 from M1. Population neural trajectories in the preparatory subspace of PMd (but not M1) were progressively less expansive with higher levels of uncertainty. Despite differences in preparatory states associated with uncertainty, the movements produced were the same. The narrower separation between states suggested a neural-based explanation for more rapid movement re-preparation with higher uncertainty. Furthermore, we found a dimension in neural state-space representing the level of uncertainty during movement preparation and execution.

**SIGNIFICANCE STATEMENT:** The dorsal premotor cortex (PMd) in primates plays a critical role in organizing preparatory states needed for the execution of target-directed movements mediated downstream by the primary motor cortex (M1). But what if the location of the target is uncertain - as often occurs in our daily lives, such as reaching for a pair of glasses in the dark that have fallen to the floor. We found that different levels of uncertainty are clearly represented in population neural activity during preparation for movement in monkey PMd but not M1. Furthermore, these different neural states related to uncertainty were not associated with changes in the movements produced. These findings provide further insight (and questions) about the operations of the motor cortex.

## INTRODUCTION

Voluntary movements are intentional actions that are cortically driven. Substantial evidence suggests that activity in the primate dorsal premotor cortex (PMd) is set up in advance of the cortical activity that instigates voluntary movements towards an external target (Weinrich & Wise 1982; Weinrich et al. 1984; Riehle & Requin, 1989; Mushiake et al. 1991; Crammond & Kalaska, 1994; Shen & Alexander 1997, Cisek & Kalaska, 2002). Such preparatory activity is thought to establish distinct neural states seeding impending neural dynamics that dictate execution of specific movements (Churchland et al. 2006a, 2006b; 2010; 2012; Ames et al. 2014; Churchland and Cunningham, 2014; Hennequin et al. 2014; Sussillo et al. 2015; Elsayed et al. 2016; Michaels et al. 2016; Lara et al. 2018; Gallego et al. 2018; 2020).

In many situations, voluntary movements are prepared and executed with incomplete information about a movement goal. For instance, reaching to a bedside light-switch in the dark depends on imprecise memory of the switch location. Likewise, when one attempts to catch a ball, one needs to *predict* (given the relative slowness in visual processing) where the ball will be upon entering reach space. On the other hand, reaching out to grab a ball that sits on a table involves minimal uncertainty. Such differences in uncertainty likely alters the characteristics of the sensory (or memory) signals that shape preparatory activity. Indeed, uncertainty-related changes in the activities of individual PMd neurons have been observed (Cisek & Kalaska 2005; Dekleva et al. 2016; 2018; Glaser et al. 2018; Suriya-Arunroj & Gail 2019). Here we explored how uncertainty about movement goals manifest in the evolution and destination of preparatory dynamics in neural populations, even when the imminent movements are the same.

We found that preparatory activity in the non-human primate PMd, but not primary motor cortex (M1), was delayed and less expansive under higher levels of uncertainty when a monkey reached to the same targets. Interestingly, this narrower preparatory space offers a neural-based explanation for more rapid re-preparation of movements when targets abruptly change location under conditions of higher uncertainty. Additionally, different levels of uncertainty were clearly represented in neural state space throughout movement preparation and execution. This indicates that features of movement *context*, such as the degree of uncertainty about a movement goal, are discernable in neural population activity of the motor cortex.

## METHODS AND MATERIALS

### Subject and Surgery

A seven-year-old (13 kg) adult male macaque monkey (*Macaca mulatta*) was trained to reach to targets for juice rewards. Approval for the animal protocol was provided by the University of Arizona Institutional Animal Care and Use Committee (IACUC) and compliance with the United States Public Health Service policy, as outlined by the Office of Laboratory Animal Welfare (OLAW), was maintained throughout the experiments. The monkey was trained to achieve proficiency in a center-out reach task described below. Following the preliminary training, we performed a sterile surgery to implant a chamber on the monkey’s skull that provided access to the premotor cortex (PMd) and primary motor cortex (M1). Additional training was required after the surgery for a total of ∼ 12 months of training. Once task proficiency was achieved, a craniotomy was performed exposing the intact dura mater in the left hemisphere directly above the PMd and M1.

### Behavioral Task

The monkey was trained on a novel variant of the forced reaction time task (Schouten & Bekker 1967, Haith & Krakauer 2015), also called the Ready-Set-Go task (Remington et al. 2018). The monkey was seated in a chair, 60 cm in front of a computer screen. With the left arm restrained, the monkey performed center-out reaching movements to two diametrically opposed targets by using his right arm and hand to manipulate a 30-cm joystick that controlled a cursor on the computer screen. Joystick signals were translated into cursor positions using NIMH Monkey Logic and MATLAB. To initiate a trial, the monkey had to voluntarily move a green cursor to a central gray circle on the computer screen (Fig. 1a, left box). A key aspect of this task was the requirement for the subject to initiate a movement almost simultaneously with a go-cue. A sequence of three timing cues (the 3^rd^ cue being the go-cue), all equally spaced in time (500 ms apart), were presented to the subject. These timing cues (flashed white rings in the center of the screen) enabled the animal to predict the occurrence of the go-cue, thereby allowing precise movement initiation.

**Figure 1.**
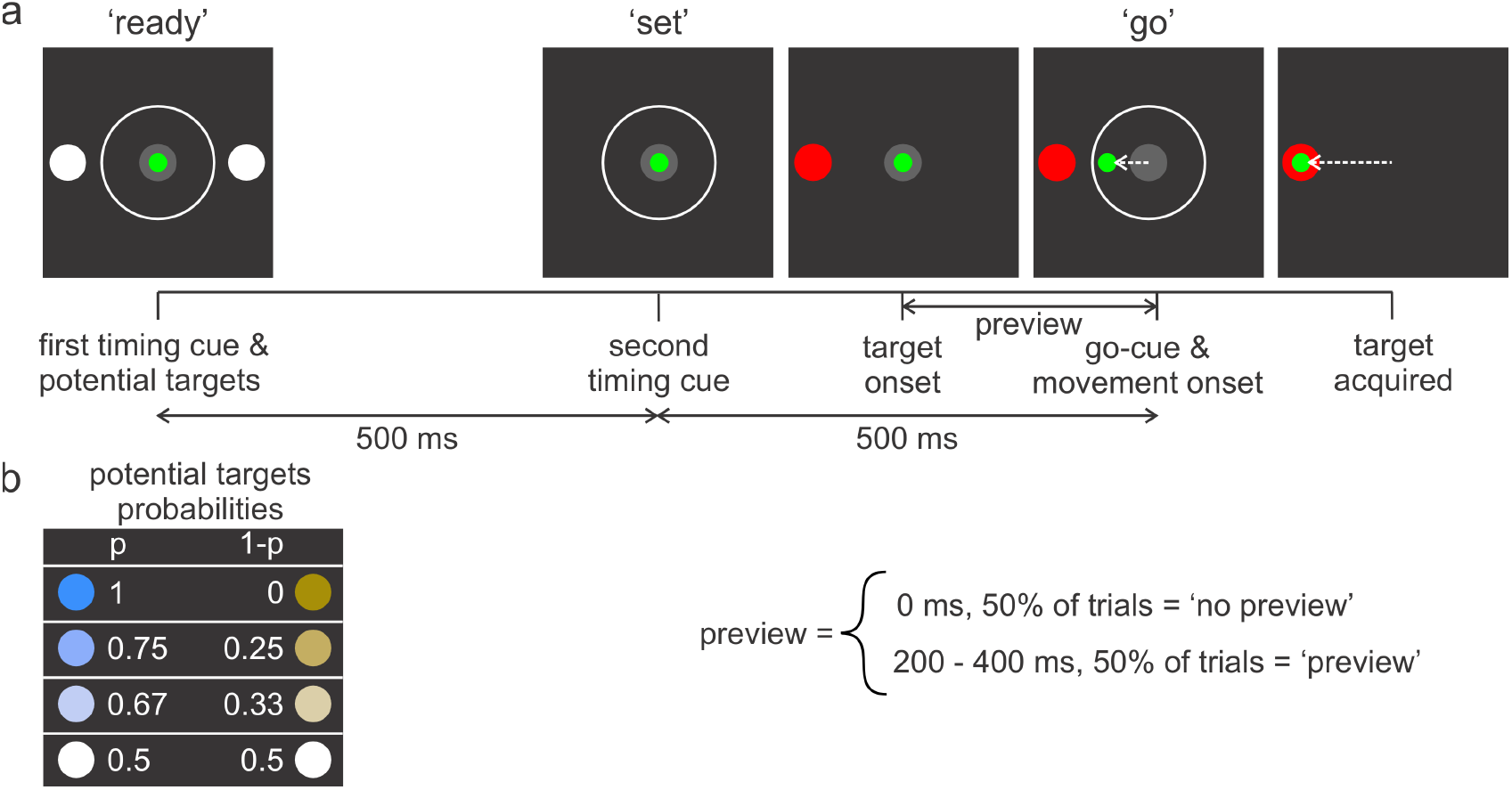
a) A Ready-Set-Go task was used to control movement onset and manipulate the level of uncertainty. When the monkey moved a green cursor using a long joystick within the center gray circle displayed on a computer screen, two potential targets were displayed along with the first timing cue (white ring – ‘ready’). 500-ms later, a second timing cue was displayed alone (‘set’). At varying times (200 – 400 ms) prior to the go cue (third flash of white ring), the actual target location (red circle) was displayed to the animal. In half the trials, no such preview was provided (i.e., the target and go cue were displayed simultaneously). The animal was required to initiate movement simultaneously with display of the third flash of the white ring (‘go’). On no-preview trials, the animal needed to guess where the target would be based on probability clues provided by the colorings of the potential targets. b) Colors and intensities of potential targets indicated the associated probability (p) of the final target location. For example, when both potential targets were white, there was equal likelihood that the actual target would be on the left or right side.

The two potential targets were displayed at the time of the first flash (Fig. 1a, ‘ready’, left box) and then disappeared coincident with the second flash (Fig. 1a, ‘set’, 2^nd^ box). The actual target (red circle 10 cm away from the center) was then displayed at varying times (Fig. 1a, 3^rd^ box) prior to ‘forced’-movement initiation at the go-cue (Fig. 1a, ‘go’, 4^th^ box). On half the trials, the time between target onset and go-cue was 0 ms (‘no preview’), while the delays for the rest of the trials were sampled from a uniform distribution that ranged from 200 to 400 ms (‘preview’). No-preview trials required the animal to initiate a movement based on his prediction of the target location. Preview trials, for which the time between target onset to movement onset was > 200 ms, enabled the animal to prepare his movement based on the visual feedback of the target. Preview and no-preview trails were interleaved in random sequence.

We manipulated uncertainty about the target location using a color scheme that indicated the probability (p) of where the final target would appear. A greater value of p, therefore, corresponded to a lower level of uncertainty. We used four different paired values of p (Fig. 1b): p = 0.5 for both targets (white), indicating the final target could appear at either location with equal probability, p = 0.66 (light blue) and 0.34 (light gold), p = 0.75 (darker blue) and 0.25 (darker gold), and p = 1 (dark blue) and 0 (dark gold). The positions (i.e., left or right) of the high/low probability targets were varied randomly across trials.

Experimental sessions consisted of several blocks of 80 successful trials. A successful trial was one in which movement initiation occurred within 100 ms of the third flash (go-cue) and the cursor intersected the target within 1000 ms of the onset of the go-cue. Position signals acquired from the joystick were low pass filtered (cutoff frequency = 4 Hz) and differentiated to determine the velocity of the joystick (and therefore the hand). Movement onset was determined as the first time the speed of the hand exceeded 5 cm/s. For each successful trial, the monkey received a juice reward and auditory feedback (chime). Failure to accomplish the task within the limits of the timing criteria resulted in automatic cessation of the trial without reward. Each block was divided into four segments; the first segment displayed the high probability potential targets, and with each subsequent segment of 20 trials, the level of uncertainty was increased in accord with the target coloring as outlined above. Such segmentation facilitated learning of probabilities by the monkey. Typically, the monkey completed a minimum of 6 blocks (of 80 successful trials) in each session.

### Neural Recordings

We performed recordings from PMd and M1, identified using structural magnetic resonance imaging scans that were aligned with a 3D model of the chamber. In each session, a sharp guide-tube passed through an opening of a chamber grid (openings every 750 µm) and penetrated the dura mater at designated locations in the chamber. A 64-channel microelectrode (S-probe; Plexon, Inc.), comprised of two adjacent columns of 32 recording sites (75 µm inter-column spacing, 100 µm interelectrode spacing) was passed through the guide-tube and into the cortex. Voltage signals from each channel were recorded using the Plexon data acquisition system at a sampling rate of 40 kHz. Single units were identified using an ofline spike sorter (Kilosort, Pachitariu et al. 2016).

### Neural Data Analysis

For each well isolated neuron, we aligned the spike times to various task related events, including: potential targets on, potential targets off, final target on, and movement onset. Following these alignments, we calculated the average spike count across trials in 1 ms time bins to generate peri-stimulus time histograms (PSTHs). We then convolved these average spike counts with a 40 ms Gaussian kernel to yield smooth firing rate traces.

#### Preparatory and Movement Subspaces

In order to decompose the neural state-space into preparatory and movement subspaces, we performed two pre-processing steps as described in previous studies (Churchland et al. 2012, Elsayed et al. 2016; Remington et al. 2018, Zimnik & Churchland 2021). First, we soft-normalized the firing rate of each neuron by a factor equal to the range of the firing rate plus 5 spikes/s. This step ensured that the low dimensional subspaces were not skewed by the neural responses of a few high firing rate neurons and also prevented over-inflating the contribution of low firing rate neurons. Second, we mean centered the firing rate of each neuron such that the mean firing rate across the two reach directions was zero at every time point for both no preview and preview trials. This step forces dimensionality reduction to capture reach-direction related population responses rather than condition-independent signals.

The animal was allowed to make a reach correction to acquire the final target. For no-preview trials at higher levels of uncertainty, the incidence of such movement corrections was high. Hence, to obtain sufficient trials of reaches in both directions per condition, we calculated a heading angle as the movement angle relative to the horizontal axis that intersected two potential targets 50 ms following movement onset. We used trials for which the heading angle was within ± 45 degrees relative to one of the targets, even if there was a subsequent movement correction. As such, we only included neural activity from the first segment of the movement (i.e., before movement correction). Also, for all uncertainty conditions except for the p=0.5 condition, we only used movements that were directed towards the high-probability location for neural data analysis.

Following the two pre-processing steps, we used the dimensionality reduction method developed by Elsayed et al. (2016). To identify the two subspaces, we divided neural activity into a preparatory epoch (500-150 ms before movement onset) and a movement epoch (50 ms before to 300 ms after movement onset). The timing and durations of these two epochs were based on qualitative inspection of single unit responses.

By optimizing the following objective function, we obtained a set of orthogonal bases, *V*_*prep*_ and *V*_*move*_ describing the preparatory and movement subspaces respectively:

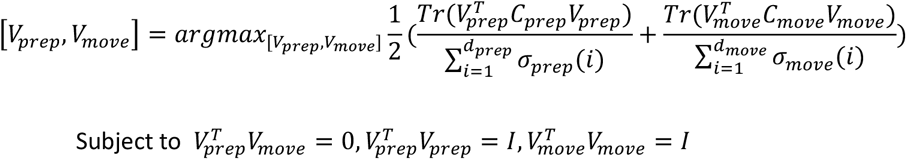

*C*_*prep*_ and *C*_*move*_ are the covariance matrices of the neural population activity during the preparatory and movement epochs respectively. *σ*_*prep*_ *(i)* and *σ*_*move*_*(i)* are the *ith* singular values of *C*_*prep*_ and *C*_*move*_ respectively. We chose the number of dimensions for both preparatory (*d*_*prep*_) and movement (*d*_*move*_) subspaces to be 10. A large percentage of reach-direction related variance was captured by the first preparatory dimension (∼75%) during the preparatory epoch and the top two movement dimensions (∼80%) during the movement epoch. The final results and conclusions were robust with respect to the choice of dimensionality. The subspace identification method described above was used for no preview and preview trials separately.

#### Uncertainty Dimension

To identify the dimension that best captured the level of uncertainty in neural state-space, we only used the no-preview trials. This was done because uncertainty about the target location was present throughout the preparatory epoch and in most cases far into the movement epoch. We started by applying the pre-processing steps similar to that mentioned in the above section i.e., first soft normalizing the firing rates followed by mean centering at every time instant. However, instead of mean centering the firing rate of each individual neuron across the two reach directions, we mean centered them across the different levels of uncertainty. Hence, the mean firing rate at every time point across the different levels of uncertainty was zero. We then applied PCA on a matrix comprising these mean centered firing rates.

#### Bootstrapping to Compare Neural Trajectories

We used the approach used by Ames et al. (2014) to determine if differences seen between neural trajectories for the different uncertainty conditions were greater than that expected by chance. For each condition (i.e., reach direction and uncertainty level), we resampled spikes times 1,000 times with replacement. The size of each resampled set was the same as the original data. We then calculated the smooth firing rate traces for the resampled data. We projected the resampled data onto the preparatory and movement subspaces. Like Ames et al. 2014, we then calculated the minimum neural distance between the resampled trajectories across the different levels of uncertainty in the preparatory and movement subspace.

To calculate the minimum neural distance, we start by choosing a reference trajectory, which is a neural trajectory for a particular reach direction and uncertainty condition. We then calculated the minimum Euclidean distance between a neural state (at one time instant) on the reference trajectory and all other neural states (at all other time instants) on the non-reference trajectory. The non-reference trajectory was chosen from a condition where the reach direction was the same as the reference trajectory, but the level of uncertainty was different. By calculating this metric between the resampled data, we were able to estimate the variability in the minimum neural distance.

To create a null distribution of minimum neural distances, we resampled trials from the same condition twice (for example, two 1,000-trial resamples for reaches made to the right under the p=1 uncertainty level) and calculated the minimum neural distance between them. To determine the likelihood that the observed neural distances were greater than chance, we calculated the probability of the across condition resampled distance being greater than the within condition resampled distance (Ames et al., 2014).

## RESULTS

### Effects of Uncertainty on Behavior

We compared hand kinematics for straight (uncorrected) reaches to targets associated with different uncertainty conditions. Kinematics were similar across levels of uncertainty (Fig. 2a). To determine whether the monkey understood the meaning of color intensity of the potential targets (and didn’t simply reach to the blue target when displayed), we calculated the proportion of trials initiated toward the lower probability (gold) target for no-preview trials.

**Figure 2.**
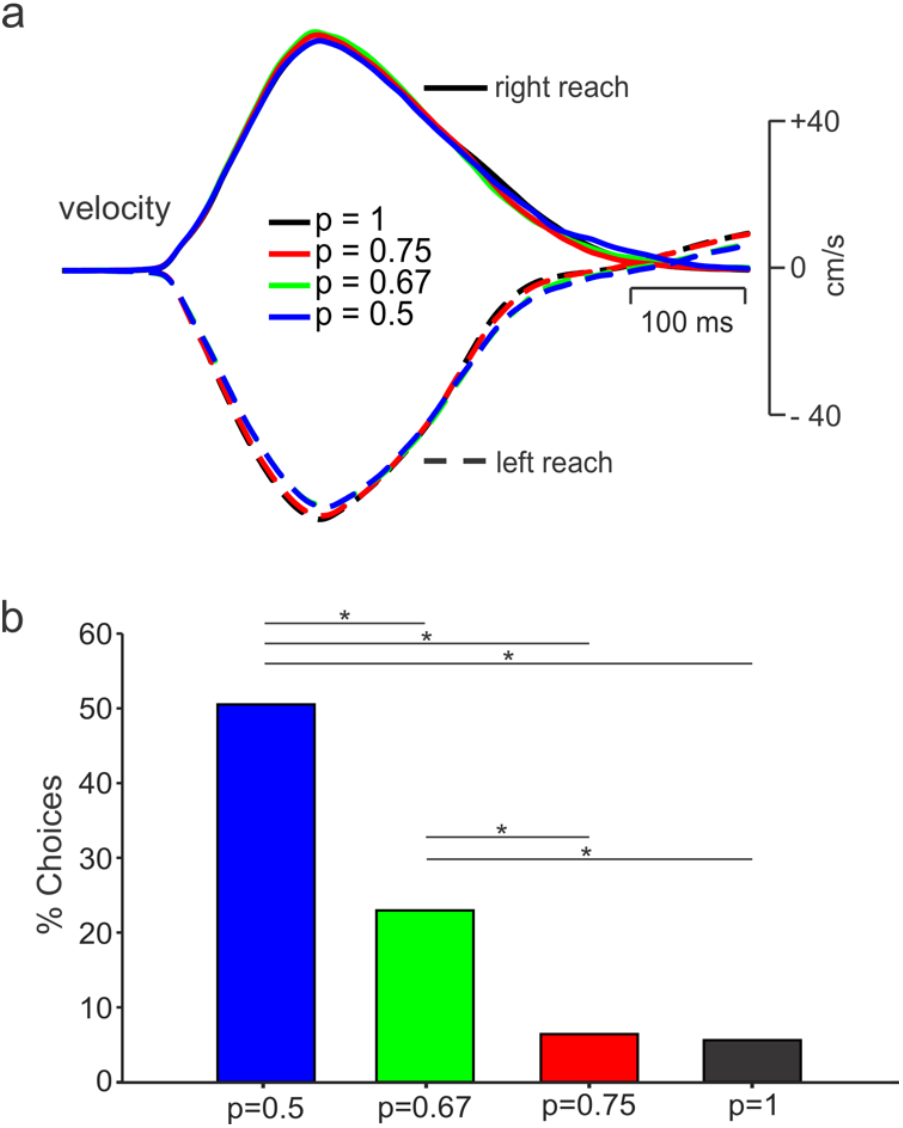
a) Average velocities of the hand for uncorrected left and right reaches during the no-preview trials across all recording sessions for the p=1, p=0.75, p=0.67 and p=0.5 conditions, respectively). b) Percentage of no-preview trials when the subject initiated movement towards the low probability (potential target) location. For the p = 0.5 condition, all no-preview trials toward the wrong target were used. (^*^ - p < 0.01, Chi-Square Test)

We speculated that with greater levels of uncertainty, the percentage of reaches toward the lower probability target should increase, as seen in previous probability matching behaviors (Saldana et al., 2022, Koehler & James, 2014). Indeed, there was a significant increase in percentage of reaches directed towards the lower probability target with increasing levels of uncertainty (Fig. 2b).

### Basic Responses of Neurons in PMd and M1

We recorded a total of 202 neurons, 165 from PMd and 37 from primary motor cortex (M1) across eight sessions while the animal performed the task (see Suppl. Fig. 1 for recording locations). Peri-stimulus time histograms (PSTHs), aligned to movement onset, were generated for each neuron. Example PSTHs for two PMd neurons are shown in Figs. 3a and 3b. These neurons showed modulation based on the level of uncertainty (colored lines) and reach direction (dashed vs. solid lines). Over the population, however, responses were highly heterogenous and could be complex, as reported previously (e.g., Churchland et al. 2010; Churchland et al. 2012; Sussillo et al. 2015). Figs. 3c and 3d show example PSTHs for two M1 neurons. While these neurons exhibited different patterns of activity for right versus left movements, there was little difference across uncertainty conditions. Furthermore, onset of activity was delayed compared to PMd. These features were generally consistent across the M1 neurons recorded.

**Figure 3.**
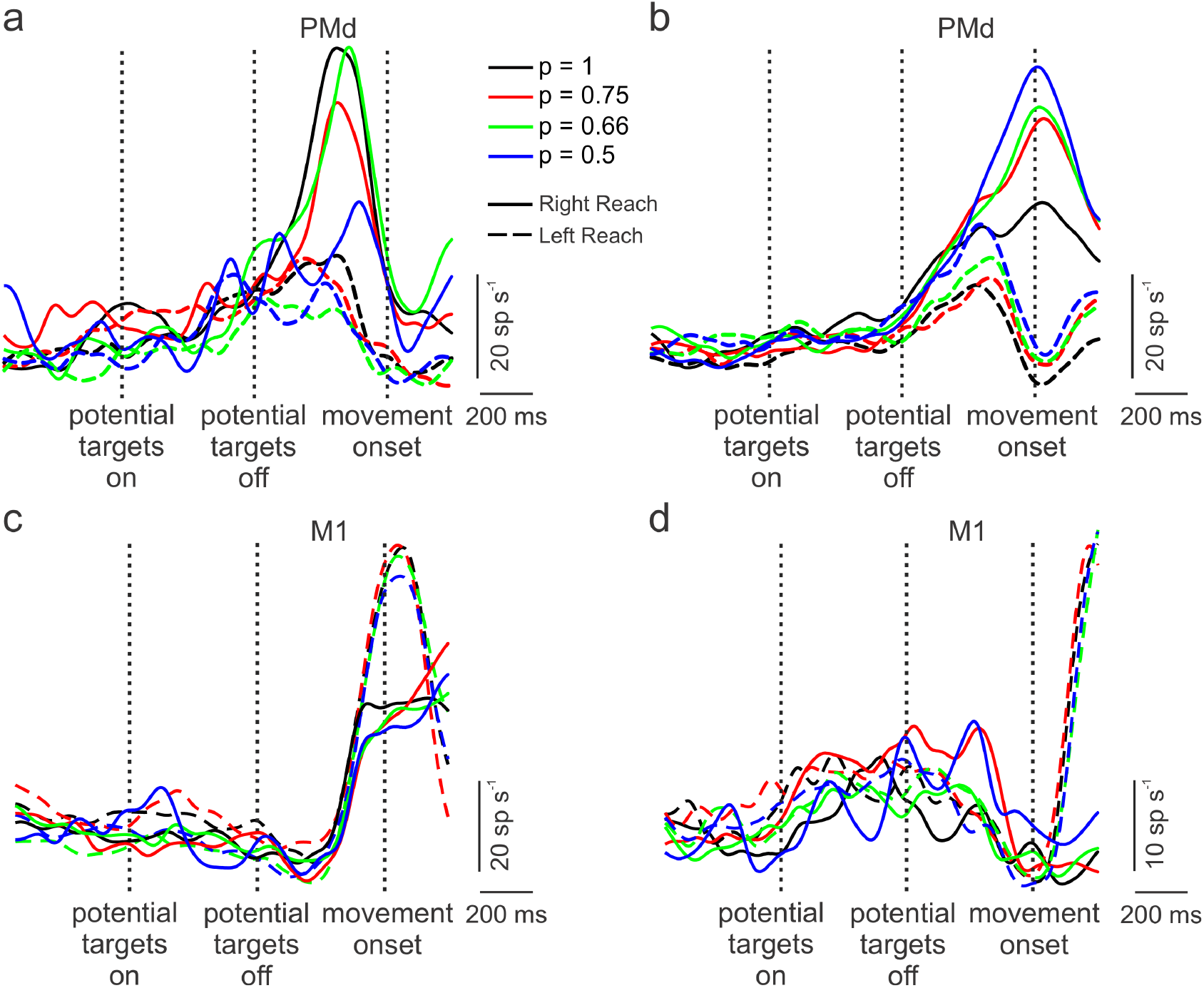
Example peri-event time histograms, aligned to movement onset, for single neurons recorded in dorsal pre-motor cortex (PMd: a, b) and primary motor cortex (M1: c, d) for no-preview trials during ready-set-go task to left (dashed lines) or right (solid lines) targets. The probabilities that a target would appear at a given location, based on the coloring of potential targets, are shown as different colored lines. Neural activity was generally different for right and left movements for both the PMd and M1 neurons. For PMd neurons, changes in neural activity occurred about 500 ms prior to movement onset whereas increased neural activity occurred later for M1 neurons. The PMd neurons show some segregation in neural activity associated with the level of uncertainty but this was not the case for the M1 neurons.

### Reduction in Preparatory Subspace Occupancy with Increasing Levels of Uncertainty in PMd

We examined the collective behaviors of all neurons recorded in PMd and M1 separately using a neural dynamics approach (see reviews by Shenoy et al. 2013, Churchland & Shenoy 2024), wherein the firing rates of each neuron represented a different dimension in neural state space. We used a dimensionality reduction technique introduced by Elsayed et al. (2016) to decompose the high dimensional neural state-space into orthogonal *preparatory* and *movement* subspaces. The time course of the first preparatory dimension (which captured ∼ 80% of the reach-direction related variance) for the PMd neural population is shown for *no-preview* trials in Fig. 4a. Preparatory activity not only was clearly different for left versus right movements but also distinguished different levels of uncertainty. The most expansive preparatory activity was observed for the most certain condition (black lines, Fig. 4a) and was progressively less with greater uncertainty (red, green, blue lines). Bootstrapped resampling (see Methods and Materials) was used to establish 95% confidence intervals for each preparatory trajectory (see Suppl. Fig 2.). The distance in neural state space between pairs of uncertainty conditions at the peaks of preparatory activity were calculated for left and right reaches separately. These distances were all significantly (p < 0.05) greater than that expected due to chance except for uncertainty conditions between the p = 0.66 and p = 0.5 for left reaches (see Suppl Fig. 2 and methods for details).

**Figure 4.**
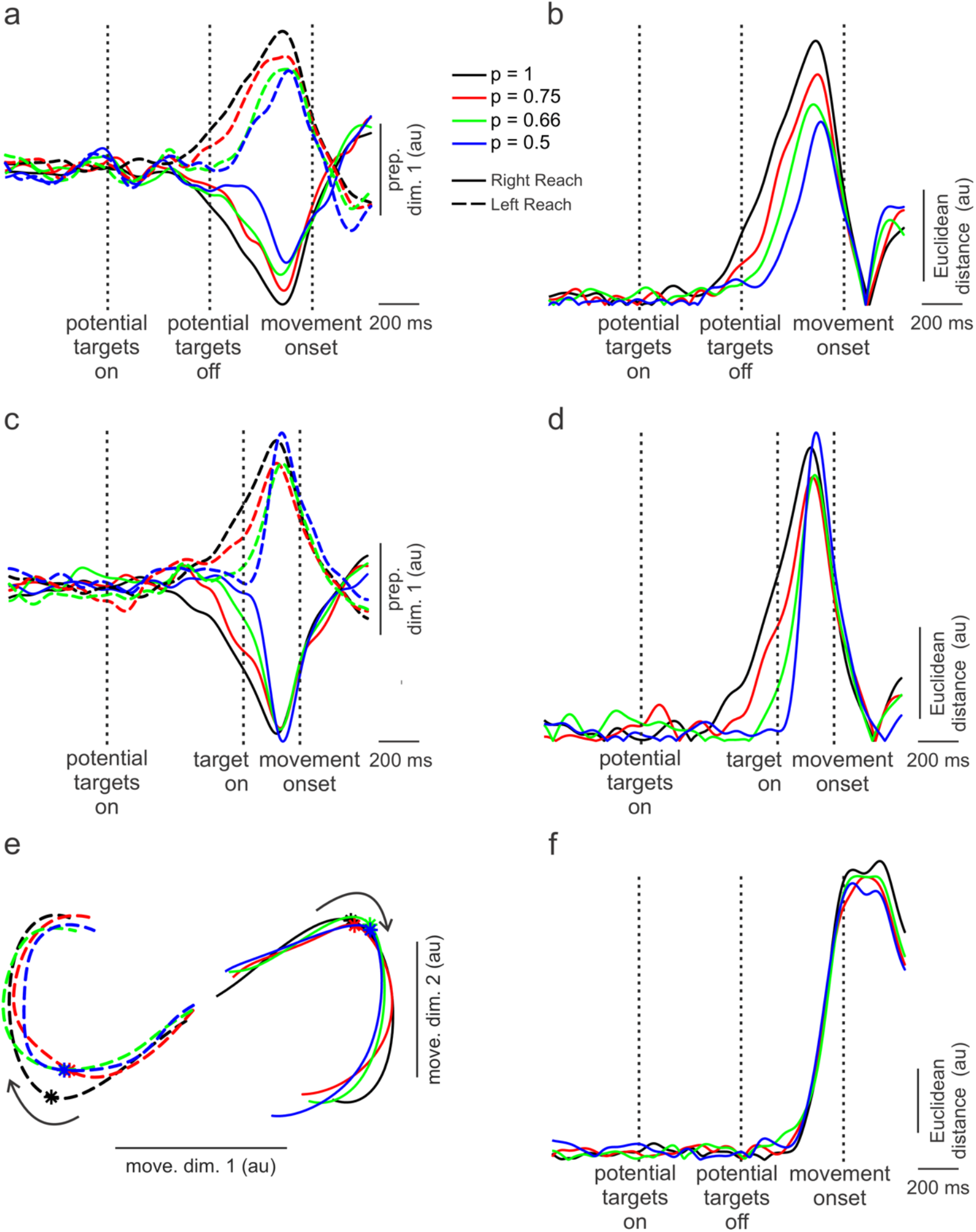
Population trajectories in preparatory and movement dimensions of neural state space in dorsal premotor cortex. a) Time courses of the first preparatory dimension (accounting for >80% of variance) for no-preview trials. Preparatory activity distinguishes both right and left movements and levels of uncertainty. b) Euclidian distances in neural state space between left and right movements. Such Euclidian distances were smaller for trials involving the higher uncertainty. c) Same as for a) but for the preview trials. In this case, separation in the preparatory dimension across levels of uncertainty collapsed as ample time of target preview (200 – 400 ms before go cue) greatly curtailed uncertainty. d) like in b) but for preview trials. Little separation in Euclidian distance between right and left movements across different levels of uncertainty at peak preparatory activity. e) Trajectories of neural activity in two movement dimensions (accounting for > 80% of the variance) in state space for left and right reaches and across levels of uncertainty. These trajectories start 200 ms before and end 150 ms after movement onset. Curved arrows show progression of time and asterisks denote onset of movements. As expected, left and right movements were clearly delineated but there was little separation across levels of uncertainty. f) Euclidian distances as a function of time in the movement subspace between left and right movements. There was little difference in this distance across levels of uncertainty.

To better illustrate these differences, we calculated the Euclidian distance in neural state space between right and left movements for each level of uncertainty (Fig. 4b). The reduction in Euclidian distance (‘occupancy’, Zimnik & Churchland 2021) with increasing uncertainty was accompanied by a delayed emergence of preparatory activity (∼300 ms later for the most uncertain compared to the most certain conditions). This might imply that with greater uncertainty (discerned by the monkey from potential-target coloring), PMd activity was held in abeyance to enable possible additional visual feedback of the target location.

### Convergence of Preparatory States during Preview Trials in PMd

The differences in preparatory states across levels of uncertainty occurred while the animal prepared movements based on *predictions* of target locations (*no-preview* trials). However, on *preview* trials, the animal had sufficient time (200 – 400 ms) just prior to forced-movement onset to prepare a movement based on visual feedback of the target. Up until the moment of target display, though, the animal presumably relied on coloring of potential targets displayed early on in the trial to ascertain likelihoods of final target locations. Consequently, at the outset of the *preview* trials, preparatory activity might be comparable to that of *no-preview* trials with different levels of uncertainty represented. Later, however, after target display, there would be little ambiguity about target location and preparatory activity should converge to a single state associated with the lowest level of uncertainty. Indeed, this is what appeared to be the case. Fig. 4c shows preparatory activity for PMd neurons during the preview trials. At target onset, there was substantial separation in preparatory activity across levels of uncertainty. Within about 150 ms of target display, however, preparatory activity converged. Even activity associated with the most uncertain condition (Fig. 4c, p = 0.5, blue traces) intersected that of the least uncertain condition (p =1.0, black traces). Such convergence is highlighted in similar Euclidian distances measured near peak preparatory activity between left and right movements (Fig. 4d).

### Little Distinction of Uncertainty in the Movement Subspace in PMd

As outlined in the Introduction, a number of studies have posited that preparatory states are necessary pre-cursors to subsequent movement dynamics. Thus, we next investigated whether different preparatory states associated with different levels of uncertainty preceded distinct movement subspace dynamics. In this case, two dimensions were needed to capture > 80% of the variance in the movement subspace. As shown in Fig. 4e, PMd activity in this 2D space was clearly different for left versus right reaches but there was little distinction across uncertainty conditions, as evinced by practically equivalent Euclidian distances between neural states for left and right movements across levels of uncertainty (Fig. 4f). Therefore, despite clear differences in the preparatory states for varying levels of uncertainty, these differences did not manifest in movement subspace dynamics in PMd, nor in the actual movements produced (Fig. 2a).

### Preparatory and Movement Subspace Dynamics in M1

Preparatory and movement subspace dynamics for M1 neurons are shown in Fig. 5. While left and right movements are distinguishable in the preparatory subspace of M1 (Fig. 5a), separation across levels of uncertainty are modest. Accordingly, Euclidian distances between left and right reaches in the preparatory subspace (Fig. 5b) were small, with only the least uncertain (p = 1, black trace) condition exhibiting slightly greater occupancy than the other uncertainty conditions.

**Figure 5.**
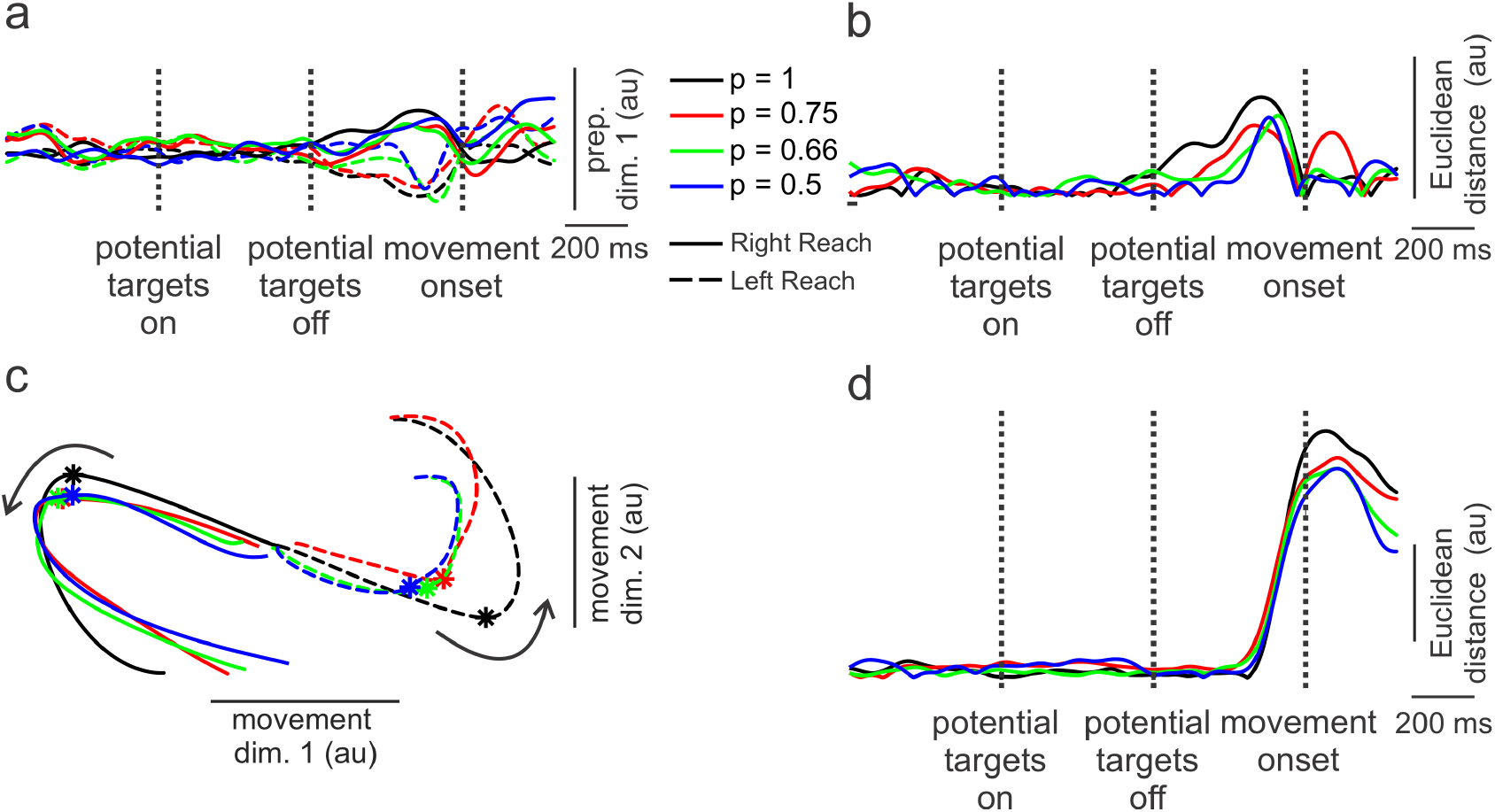
Population trajectories in preparatory and movement dimensions of neural state space in primary motor cortex (M1). a) Time courses of the first preparatory dimension subtly distinguish right and left movements but not levels of uncertainty. b) Euclidian distances in neural state space between left and right movements exhibited little separation across levels of uncertainty. c) Trajectories of neural activity in two movement dimensions for left and right reaches and across levels of uncertainty. Curved arrows show progression of time and asterisks denote onset of movements. Like PMd (Fig. 4e), left and right movements were clearly delineated but there was little separation across levels of uncertainty. d) Euclidian distances in neural state space as a function of time in the movement subspace between left and right movements.

Like for PMd, two dimensions of M1 neural dynamics were needed to capture > 80% of the variance in the movement subspace (Fig 5c). Also, as was the case for PMd, the two directions were clearly distinguished in the movement subspace. In addition, there was minimal separation across levels of uncertainty in this subspace (like PMd), highlighted in the similarity and extent of the Euclidian distances between left and right reaches (Fig. 5d). Only the peak Euclidian distance for the most certain condition (p = 1.0) was somewhat greater than for the other conditions (Fig. 5d). This occurred because the neural trajectory of the higher certainty condition had a greater magnitude but only for left reaches (dashed black line, Fig. 5c). The reason for this is not clear but may be related to greater variability in M1 population trajectories possessing relatively few cells compared to PMd.

### Re-preparation is faster with higher uncertainty

Given that state-space distances for preparation in PMd between left and right movements were smaller for cases with the most uncertainty (Fig. 4b), we asked whether the relative nearness of those states might enable faster movement re-preparation upon receiving visual feedback of the final target location. In other words, if initially planning to move in one direction but later information becomes available indicating that the target is in the opposite direction, then the distance to traverse in state-space to the updated preparatory state would be less under conditions of high uncertainty. This might reduce the time needed to re-prepare the movement.

We examined this possibility by quantifying the heading error (angle between the initial reach direction and final target location) as a function of the time elapsed between target onset and movement onset for no-preview and preview trials combined. We considered an ‘accurate’ reach as one in which the heading error was less than 45 degrees. Fig. 6a shows an example set of trials for the p = 0.5 condition. As expected, for brief target-onset to movement times, about half the trials were in the ‘correct’ direction to the final target (0 degree heading error) and about half were to the ‘wrong’ target (180 degree heading error), indicating the animal was guessing. Trials for which target-onset to movement time were negative are those in which the monkey ‘jumped-the-gun’ and started moving before the target was displayed. When the times between target and movement onsets were long (> ∼ 200 ms), almost all the trials were accurate.

**Figure 6.**
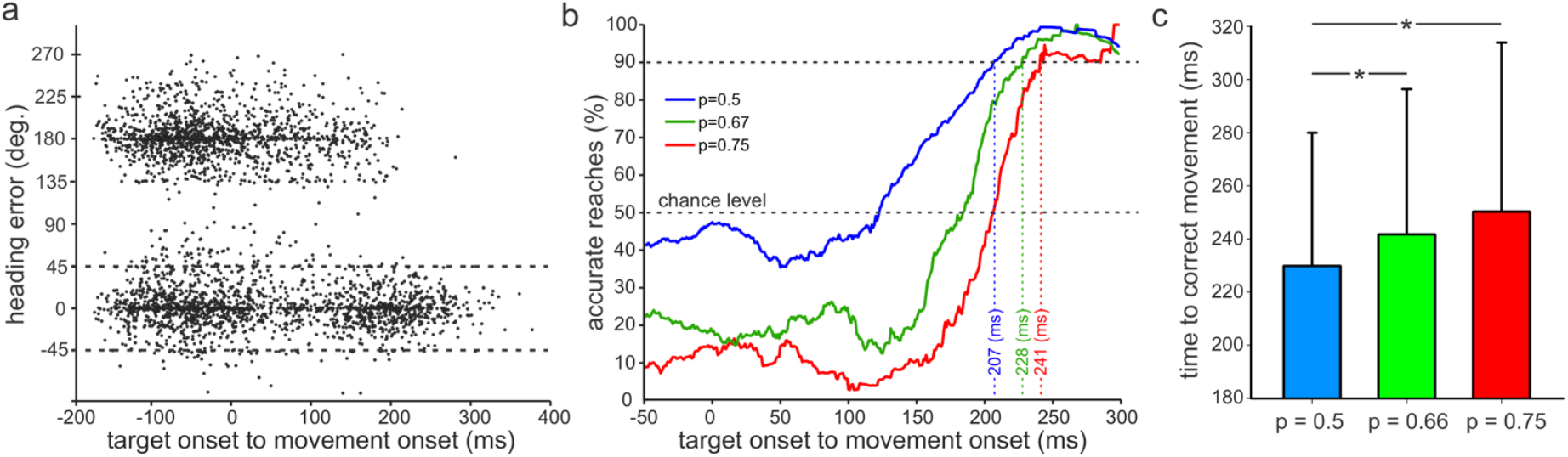
Time required for re-preparation in both no-preview and preview trials combined. a) Heading error as a function of the time from target onset to movement onset under the p = 0.5 condition. Each point corresponds to a single successful trial. The horizontal dashed lines were used as the threshold for identifying accurate reaches. Trials indicated with negative movement-onset times were those in which the animal ‘jumped the gun’ – initiating movement prior to the go cue. b) Percentage of accurate reaches in a sliding 40 ms time window for the different levels of uncertainty. The blue trace is directly derived from the data shown in a). The red (p = 0.75) and green (p = 0.67) traces are constructed using trials when the final target appeared at the **lower** probability location. We assumed that in such cases, the animal initially prepared movements to the higher probability target and then had to reprepare the movement when the low probability location appeared as the actual target. Vertical dashed lines indicate time needed from actual target display to movement onset to achieve 90% accurate reaches. This occurred earlier for trials with higher uncertainty. c) Mean (SD) time between target display and onset of the movement correction for trials when the animal altered course after initiating a movement in the wrong direction. There was a significant main effect of uncertainty on correction time (p < 0.001, Kruskal-Wallis 1-way ANOVA, n = 715, 521, 473 for the p = 0.5, 0.67, and 0.75 conditions, respectively). ^*^ - p < 0.05, Dunn’s Method post-hoc analysis.

We determined the time between target and movement onsets that was required for 90% of the reaches to be accurate (Fig. 6b). To do this, the percentage of accurate reaches (i.e., those with heading errors < ± 45 degrees) were calculated from the raw data (like that shown in Fig. 6a) in 50-ms bins across the range of preview times. We used all p = 0.5 trials in these calculations (blue line, Fig. 6b). For the p = 0.67 (green line, Fig. 6b) and p = 0.75 (red line, Fig. 6b) conditions, we selected trials for which the final target location appeared at the *lower* probability site. We did this because we were interested in those cases where a movement was likely re-prepared (i.e., from a movement prepared to the high-probability location to the low-probability location). For p = 0.5 trials, we could not ascertain trials for which re-preparation occurred. Likewise, for the p = 1.0 trials, virtually all movements were to the high probability target, with few to the low probability location (and hence is not shown).

At brief target-to-movement times, the percentage of correct reaches (Fig. 6b) reflected what would be expected for differing degrees of uncertainty, keeping in mind these data represent cases for which the final target was at the low probability location. With increasing time between the display of the target and movement onset, the percentage of accurate movements increased steeply. For example, for the least uncertain condition (p = 0.75), that steep increase began at about 170 ms and reached 90% accurate reaches at 241 ms. For the more uncertain (p = 0.67) condition, the steep rise in percentage of accurate reaches appeared to begin slightly earlier (at a target-to-movement time of ∼150 ms) and reached the 90% level at 228 ms. And for the most uncertain condition (p = 0.5), improvement in percentage of accurate reaches commenced at about 125 ms and attained the 90% level at 207 ms (Fig. 6b). The reduction of time needed to putatively re-prepare movements and accurately reach straight to the target in the lower probability (p = 0.67) compared to higher probability (p = 0.75) condition might have transpired because of the shorter ‘travel’ distance from one preparatory state to the other under conditions of higher uncertainty (Figs. 4b). Furthermore, on trials when the monkey made a course correction after launching a movement in the wrong direction, the time between target display and onset of the movement correction was briefer for the most uncertain condition (Fig. 6c).

### Persistent uncertainty dimension independent of reach direction

A number of studies have shown that neural activity in motor cortex can represent non-movement related variables, such as expected reward size and type of reward (Ramkumar et al. 2016; Ramakrishnan et al. 2017; Smoulder et al. 2024; Derosiere et al. 2025). In addition, activities of individual PMd neurons have been shown to be influenced by the extent of uncertainty during voluntary movements (Cisek & Kalaska 2005; Dekleva et al. 2016; 2018; Glaser et al. 2018; Suriya-Arunroj & Gail 2019). Along these lines, we investigated whether a dimension in neural state space exists that encodes the level of uncertainty independent of movements produced or other neural dimensions. Indeed, we found an “uncertainty dimension” that was orthogonal to the preparatory and movement dimensions in PMd. Fig 7a shows the time course of neural dynamics along the first uncertainty dimension for the *no preview* trials. Note that left and right movements were not distinguished in this dimension. Interestingly, activity in this dimension only begins to emerge 500 – 600 ms after the potential targets were displayed with color indicating the probability levels for the forthcoming target location. For *preview* trials (Fig. 7b), with 200 – 400 ms viewing time of the actual target location prior to the go cue, activity in the uncertainty dimension was markedly foreshortened.

**Figure 7.**
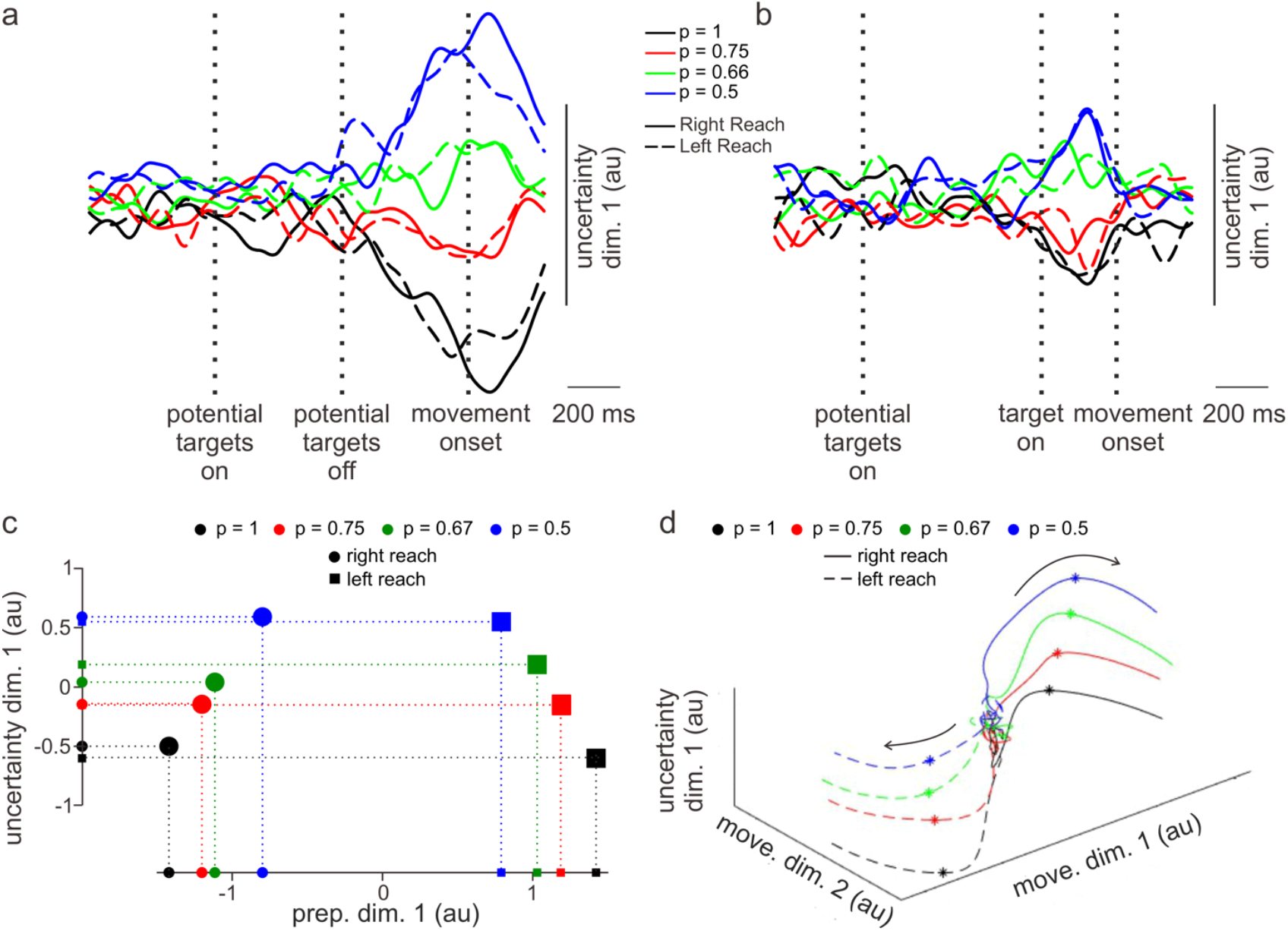
Uncertainty dimension in PMd neural state space. a) Time course of dynamics along the first uncertainty dimension (∼65% of uncertainty related variance) for no preview trials and b) for preview trials. c) Neural states along the first preparatory (x axis) and uncertainty dimensions (y axis) 175 ms before movement onset (about the time preparatory subspace activity transitions to the movement subspace). d) 3D plot with the x-y plane representing the movement subspace and the z axis representing the uncertainty dimension. Asterisks denote time of movement onset and arrows indicate progression of time.

Fig. 7c shows the single neural states associated with different movements and levels of uncertainty for the no preview trials 175 ms prior to movement onset (about the time activity emerges in the movement subspace, see Fig. 4f) plotted in a plane composed of the preparatory and uncertainty dimensions. Projection of these neural states onto the preparatory-dimension axis shows separation of movement direction and uncertainty levels (Fig. 7c). Projection onto the uncertainty dimension shows separation only in terms of uncertainty but not movement direction (as expected from Fig. 7a). When the uncertainty dimension was included in the two-dimensional movement subspace (Fig. 7d), activity related to uncertainty was present throughout the entire movement execution period. Activity along this uncertainty dimension was observed starting at about 400 ms prior to movement onset and was maximally active around movement onset (asterisks, Fig. 7d).

## DISCUSSION

We trained a monkey to reach one of two targets and to launch movements simultaneously with a go cue. On about half the trials, visual information about the target location was not available at the time of movement initiation. However, display of pre-targets (1 s before the go cue) indicated the likelihood that the actual target would be at one location or the other. We analyzed neural population activity in the dorsal pre-motor (PMd) and primary motor (M1) cortices prior to and during the initial stages of movements associated with varying levels of uncertainty by decomposing the neural state space into preparatory and movement subspaces. This analysis led to four main findings: 1) different levels of uncertainty were distinguishable in the preparatory subspace of PMd but not in M1; 2) despite different representations in PMd preparatory subspaces, there was little variation in movement subspaces (or actual movements) across levels of uncertainty in both PMd and M1; 3) the distance between preparatory subspaces for left and right movements diminished in PMd with higher levels of uncertainty; and 4) the time to reprepare a movement from one target to the other was briefer with higher uncertainty. In addition, we identified an “uncertainty dimension” in PMd orthogonal to the preparatory and movement subspaces, that appears to represent the level of uncertainty during both movement preparation and execution.

### Uncertainty influences preparatory subspace but not forthcoming movements

Preparatory subspace activity in PMd was more robust and expansive with lower levels of uncertainty (see Figs. 4 a,b). It is as though with increasing levels of uncertainty about an upcoming movement, the PMd did not vigorously embrace a particular preparatory state but instead ‘hedged its bet’ with less well-defined activity. A recent study by Michaels et al. (2025) also found a similar reduction in preparatory subspace occupancy based on the level of uncertainty regarding the direction of an external perturbation of the arm. Regardless, such fuzzier representations of the preparatory state in the present study were still sufficient to elicit accurate movements indistinguishable from those associated with little uncertainty. This finding raises interesting questions about the breadth of preparatory neural states that can set the stage for forthcoming dynamics driving a specific movement. For example, one can see in Fig. 7c the fairly large spans in the preparatory dimension (across levels of uncertainty) for left or right movements relative to the distance *between* left and right movements in this dimension.

In some respects, these finding parallels that of Lara et al. (2018). They showed distinct regions within the preparatory subspace that were linked to specific reach movements. Furthermore, those regions were preserved across different procedures used to instigate movements (i.e., delayed, self-initiated, no delay). Despite clustering of preparatory activity for the same reach under different conditions, in some cases, the expanse of the preparatory region for a given movement was quite large. This might simply reflect some degree of noisiness in estimating neural states from the relatively small subset of all neurons actually involved. On the other hand, it might also represent a basin of attractor states that are all drawn toward a single stable trajectory in state space that then mediates a particular movement (Carnevale et al. 2015).

### Smaller span between preparatory states for most uncertain conditions may facilitate brisker repreparation

The more certain the animal was of a target location, the earlier and more expansive preparatory neural activity was in the PMd for the no preview trials (Fig. 4a). The apogee of this neural activity, however, occurred at about the same time, and prior to movement onset, across all uncertainty conditions. A consequence of such uncertainty-related shaping of preparatory activity was that the state-space distance between left and right movements was smaller for the most uncertain conditions (Fig. 4b). Previous studies have shown that movement corrections are preceded by re-entry into the preparatory subspace (Ames et al., 2019). We were curious to know, therefore, whether such shorter distances between preparatory neural states might enable brisker repreparation when circumstances require an abrupt shift in intended movement from one direction to the other.

We were able to identify presumed cases of such switches from trials for which the actual target occurred at the low probability location. Our assumption was that the animal typically planned an upcoming movement based on the well-learned color-coding of probabilities for final target locations displayed at the outset of each trial (Fig. 1b). Indeed, in no preview trials, movement initiation was most often towards the *high* probability target location (Fig. 2b). The time required from target onset of the *low* probability target to initiation of accurate reaches to that target were briefer under conditions of higher uncertainty (Fig. 6b). Furthermore, when movements were incorrectly initiated toward the high probability target in these cases, the time needed to correct the movement ‘on-the-fly’ was also briefer (Fig. 6c). In some respects, this makes intuitive sense: the less committed one is to a plan, the more readily and swiftly can one change the plan. A possible neural mechanism that could account for such intuition is the narrower distance between preparatory states under situations of higher uncertainty.

Alternatively, the separation along the uncertainty dimension, by virtue of different local dynamics in neural state-space, could lead to different *speeds* of re-entry into preparatory subspaces, leading to faster movement corrections. Such context dependent changes in dynamics have been previously observed during other behaviors in dorsomedial frontal cortex (Remington et al., 2015) and M1 (Amematsro et al., 2025). Further experimental (and computational) efforts are required to help distinguish distances from rates of transition in preparatory subspaces as contributors to more rapid movement corrections with higher uncertainty.

### Uncertainty dimension

An unexpected finding of the present study was the presence in PMd of a subspace dimension that appeared to robustly represent the level of uncertainty during both movement preparation and execution (Fig. 7). This finding has some general similarities to those reported recently by Smoulder et. al (2024). They found a dimension of motor cortical activity in monkeys that was orthogonal to the preparatory subspace and represented the potential reward size for successful reaches to targets.

What might be the function or meaning of such context-related (i.e., uncertainty, reward, etc.) representations in motor cortex during planning and execution of voluntary movements? In the Smoulder et al. (2024) case, reward size was associated with different representations in the preparatory subspace that corresponded to differences in the speed and accuracy of reaches. In the present case, while uncertainty led to distinctly different representations in the preparatory subspace, it had no detectable impact on the movements produced.

It seems feasible that representations of uncertainty might be found throughout the brain - but such representations may not necessarily overtly influence specific processing or operations in various brain regions. For example, neurons in virtually every brain region recorded from in the mouse significantly modulated their firing rates just prior to movement onset (forelimb motion controlling a wheel), including in the hippocampus, primary visual areas, somatosensory cortex, periaqueductal gray, as well as motor cortex (Steinmetz et al. 2019). Steinmetz and colleagues (2019) also found wide-spread representations of how engaged the animal was in the task, including in motor cortex. Such context related activity might arise from neuromodulator release throughout the brain affecting arousal, attention, and cognitive processing (e.g., Aston-Jones & Cohen 2005; Shine et al. 2021). In addition, it could be that some context representations might merely reflect the massive interconnectivity of the brain. As a result, neural activity associated with processing and operations carried out in specific areas might nevertheless ripple throughout much of the brain. With powerful computational methods applied to large populations of neurons, such low-level ‘seismic’ signals might be detected far from their source. In this regard, it would be of interest to determine if clearly non-motor related contexts, such as the color of the screen upon which reach targets are presented, or whether or not an investigator is in the testing booth (Martin et al. 2023) during a reaching task, are identifiable in the motor cortex.

## Acknowledgements

Work supported by NIH grants NS102259 & MH121009. We are grateful to Derek O’Neill, Rujula Manjarekar, and Chloe Larkin for technical assistance.

## SUPPLEMENTAL INFORMATION

**Supplementary Figure 1.**
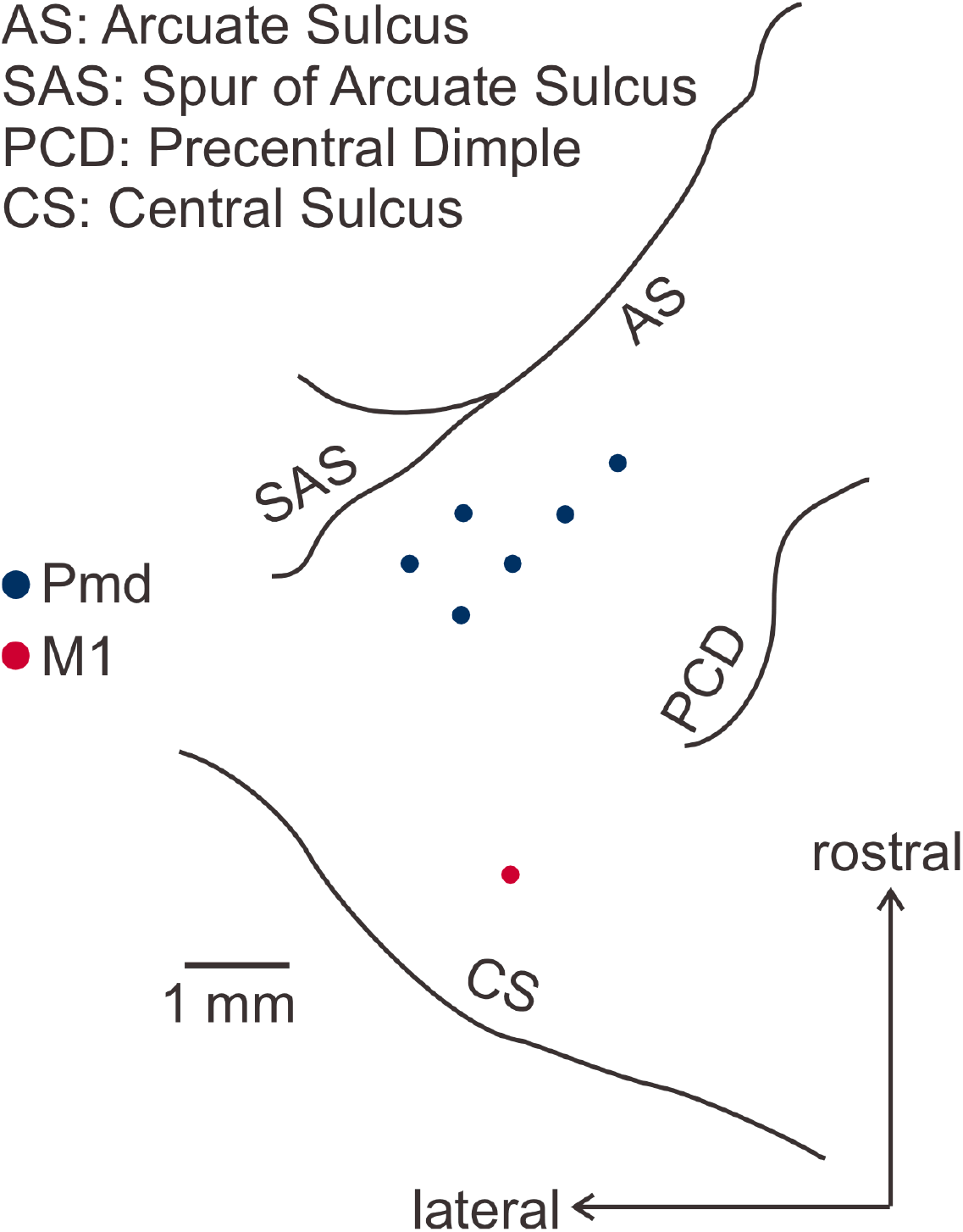
Recording sites across eight sessions. The M1 recording site was recorded from twice.

**Supplementary Figure 2.**
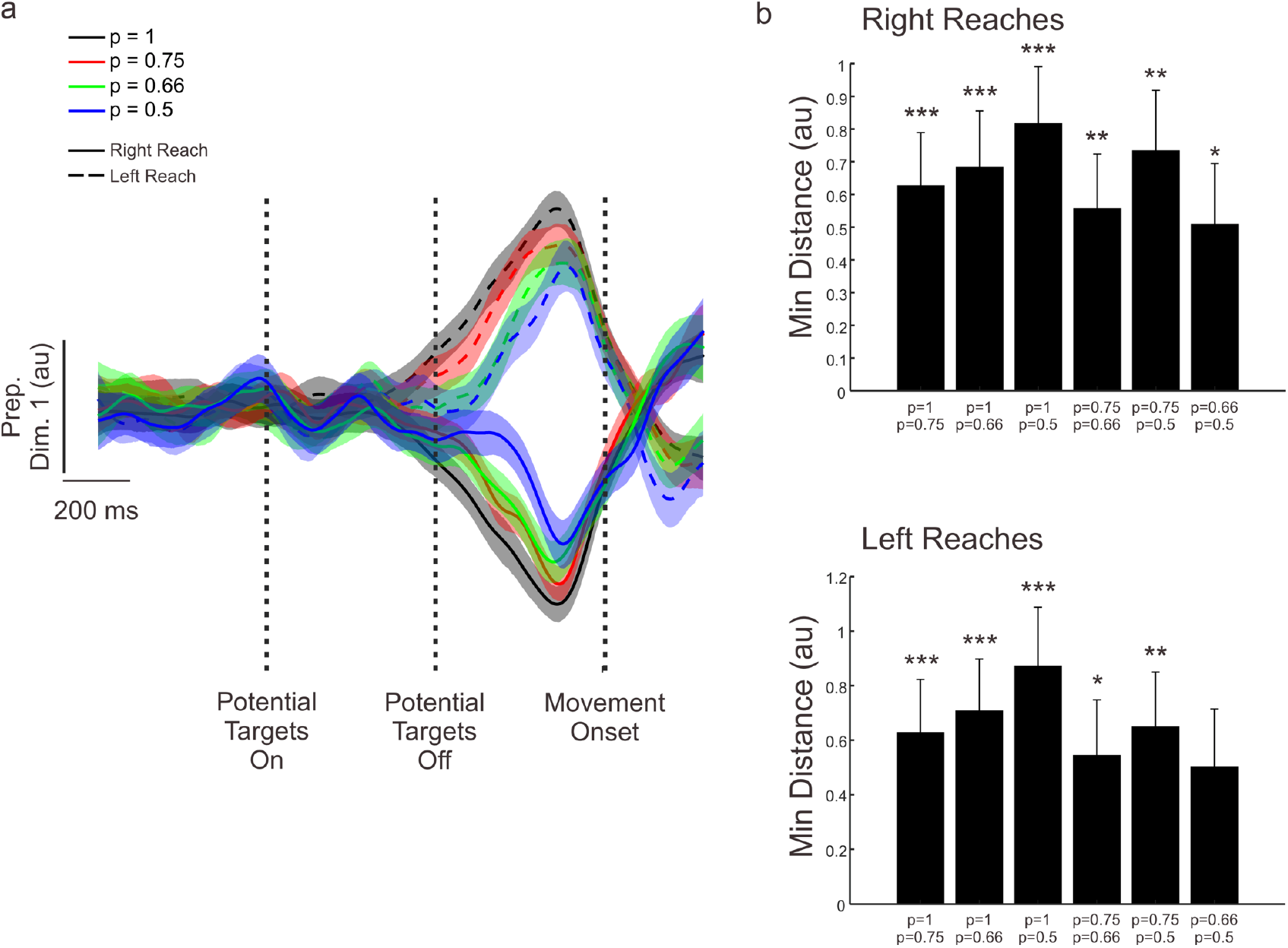
Statistical comparison of preparatory dynamics. a) Mean and 95% confidence interval of preparatory dynamics in PMd along the first dimension. b) Median minimum distance between pairs of preparatory states at peak occupancy estimated using bootstrapping. The error bars represent the 5^th^ and 95^th^ percentile of the bootstrapped minimum distances. The stars represent the bootstrap significance levels (^*^ - p < 0.05, ^**^ - p < 0.01, ^***^ - p < 0.001).

